# Mature Neurons’ sensitivity to oxidative stress is epigenetically programmed by alternative splicing and mRNA stability

**DOI:** 10.1101/2021.12.25.472549

**Authors:** Yuan Zhou, Sherif Rashad, Teiji Tominaga, Kuniyasu Niizuma

## Abstract

Neuronal differentiation is a complex process that entails extensive morphological, transcriptional, metabolic, and functional changes that dictate neuronal lineage commitment. Much less understood is the role that epigenetic and epi-transcriptional reprogramming plays in the process of neuronal differentiation and maturation. To depict the whole landscape of transcriptomics and epigenetic changes during neuronal differentiation and maturation, we differentiated SH-SY5Y cells and performed RNA sequencing on differentiated and undifferentiated cells. 728 differentially expressed genes (DEGs) enriched in synaptic signaling and cell morphogenesis pathways were observed. Moreover, transcriptome-wide mRNA stability profiling revealed that genes with altered stability were exceptionally enriched for redox homeostasis pathways. Mature neurons are known to be highly sensitive to oxidative stress, which is crucial in the pathophysiology of neurodegenerative disease. Our results suggest that this heightened sensitivity is regulated at the mRNA stability level (i.e., epigenetic) rather than at the transcriptional level. Alternative splicing analysis revealed the exon skipping and alternative mRNA isoforms enriched for morphogenesis related pathway. Alternatively, alternative 5 and 3 prime splicing site, intron retention and mutually exclusive exon events exclusively clustered in the translation and translation initiation pathways, suggesting the potential effect of alternative splicing on translation following neuronal maturation. Splice motif analysis revealed enriched motifs for RBPs that regulate various splice types and can be further correlated to distinct phenotypical changes during neuronal differentiation and maturation. Here we present an extensive exploration of the transcriptional and epigenetic changes and their potential association with the process of neuronal differentiation, providing a new insight into understanding the molecular mechanism of neuronal function and behavior.

## Introduction

Neuronal differentiation is a complex multistep process in which neurons undergo dramatic morphological alterations, including neurite outgrowth and synapse formation^1^. A series of changes then enables neuron to carry out their desired activities such as, electrophysiological activity and neurotransmitter secretion^2,3^. Consequently, these changes render mature neurons highly sensitive to oxidative stress, a property which plays a crucial etiology in many neurological and neurodegenerative diseases^4,5^. Given the important link between oxidative stress in neurons and neurodegenerative diseases such as Alzheimer’s^6^ and Parkinson’s^7^, it is crucial not only in understanding the pathophysiology of diseases, but also to develop strategies for neuroprotection via genetic or pharmacological interventions.

Recent advances have revealed an important role that the epigenome plays in regulating neuronal behavior as well as in neurogenesis. For example, a dynamic mRNA decay network regulated by RNA binding proteins (RBPs) Pumilio contributes to the relative abundance of transcripts involved in cell-fate decisions and axonogenesis during Drosophila neural development^8^. Moreover, emerging evidence unveiled a strong association between altered mRNA stability and neurodegenerative disease^9^. On the other hand, neurons at their different developmental stages have specific alternative splicing (AS) patterns, indicating a potential association of AS in regulating neuronal differentiation and maintaining neuron maturation^10^. While many post-transcriptional events occurring during the neuronal differentiation are now increasingly understood, it remains largely unknown how exactly post-transcriptional regulation contributes to neuronal differentiation. Furthermore, how the changes associated with neuronal differentiation and maturation render neurons more sensitive to oxidative stress is not fully understood.

SH-SY5Y human neuroblastoma cells are frequently used as *in vitro* neuronal differentiation model due to the biochemical and morphological resemblance of neuron to study neuronal physiology and neurodegenerative diseases^2,11^. Previous transcriptomic profiling on both differentiated and undifferentiated SH-SY5Y cells revealed that differential gene expression in these neuron-like cells are more functionally linked to changes in cell morphology including remodeling of plasma membrane and cytoskeleton, and neuritogenesis^12,13^. On the other hand, SH-SY5Y cells develop distinct responses to cellular stressors/neurotoxins after differentiation which leads to their utilization as cell model systems to study the mechanisms of neurodegenerative disorders^14^. However, existing transcriptomic analysis on differentiated SH-SY5Y cells cannot provide molecular basis to this heightened sensitivity to stress^12,13^.

The aim of this study was to elucidate the global landscape of transcriptomic and epigenetic regulatory networks involved in the process of neuronal differentiation and maturation. Using SH-SY5Y cells as a model for neuronal maturation, and utilizing RNA deep sequencing and mRNA stability analysis, we revealed that changes in the transcriptome regulate morphological and structural changes that promote neuronal function. Interestingly, mRNA stability changes, which reflect epigenetic and epi-transcriptional phenomena, revealed the basis for neuronal vulnerability to oxidative stress. This observation was corroborated with data from a recently published CRISPR screen in iPSC derived neurons^15^. Our data also revealed intense AS changes that correlate with various aspects of neuronal maturation and function. Importantly, different AS types were related to different phenomena, indicating tightly regulated and complex epigenetic reprogramming of neuronal maturation.

## Result

### Confirmation of SH-SY5Y Neuronal Differentiation

SH-SY5Y cells were subjected to 10-days differentiation treatment according to the schematic in **Figure 1A**. To verify the neuronal characteristics of fully differentiated SH-SY5Y cells, we evaluated the morphological changes of differentiated SH-SY5Y by light microscopy and the expression of mature neuronal markers by immunofluorescent staining. Undifferentiated SH-SY5Y cells grew in clumps and demonstrated a large, flat, epithelial-like cell body with numerous short processes extending outward^16^, while pre-differentiated cells by 4-day RA treatment exhibited disassociated clump, cell body shrinkage and possess several neurites outgrowth, and fully-differentiated cells by subsequent 6-day BDNF treatment developed a triangle cell body and extensive neurites that project to surrounding cells to form complex neurite network (**Figure 1B**). Moreover, **Figure 1C** demonstrated the higher expression and the polarized subcellular distribution of all the neuronal markers (Nestin, MAP-2, β-tubulin-III and Synaptophysin) in the differentiated cells (Diff.) compared to the undifferentiated cells (Undiff.), suggesting these differentiated cells have already possessed some features of neuron.

**Figure 1.**
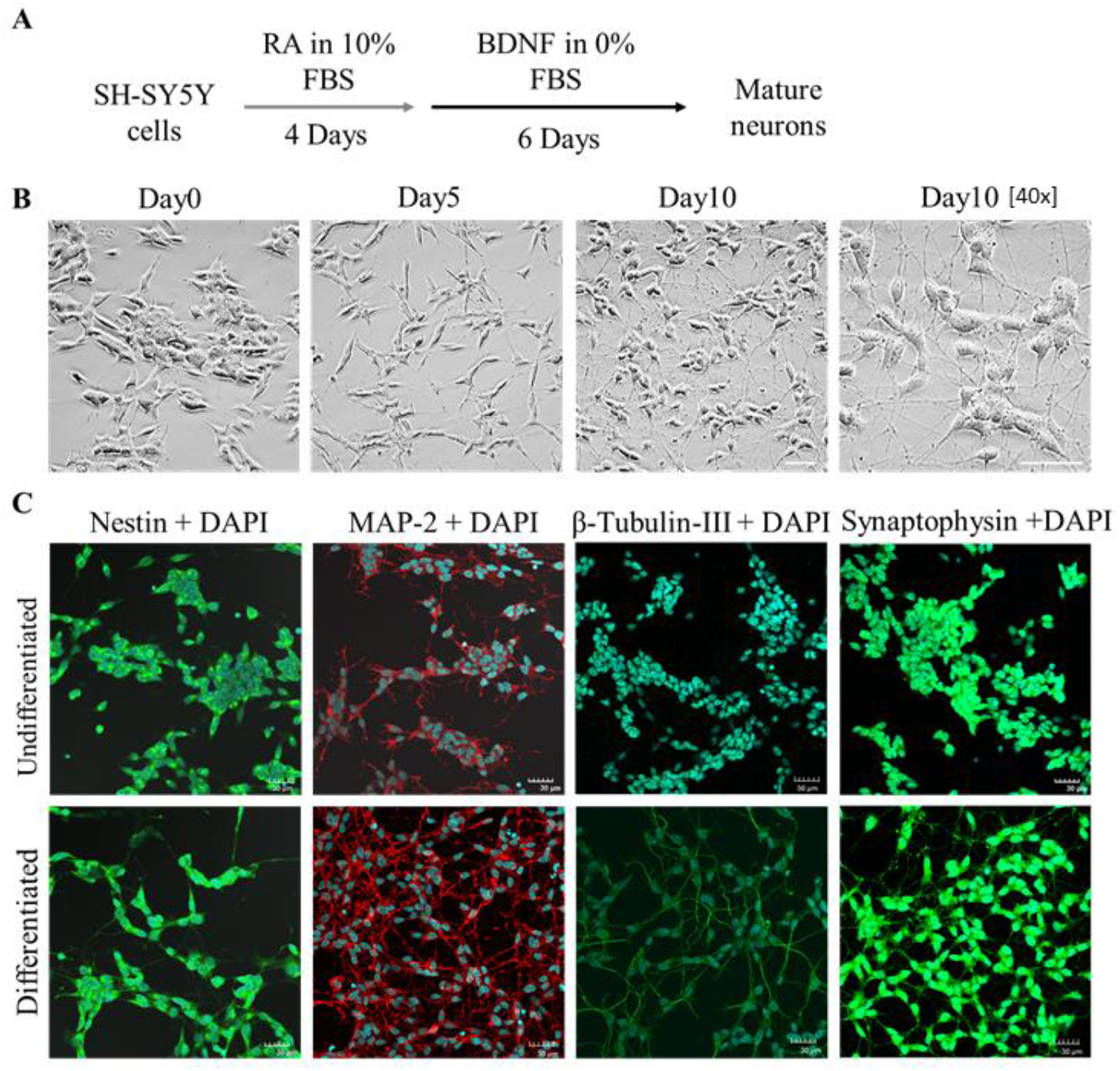
Differentiation of SH-SY5Y human neuroblastoma cells towards neuron-like cells. **A** Schematic diagram of the protocol for neuronal differentiation of SH-SY5Y cells. RA: retinoic acid; BDNF: brain derived neurotrophic factor. **B** Morphological appearance of undifferentiated (Undiff.) (Day0) and differentiated (Diff.) (Day5, 10) SH-SY5Y cells by light microscope. Images were obtained at 20X magnification and 40X magnification. Scale bar: 50μm. **C** Immunofluorescent staining on neuronal markers including Nestin (Green), MAP2 (Red), β-tubulin-III (Green), Synaptophysin (Green) with DAPI (Blue) in Undiff. vs Diff. cells at the endpoint of 10 days. Images were obtained at 40X magnification. Scale bar: 30μm.

### Transcriptomic Profiling of Differentiated SH-SY5Y Cells

Gene expression analysis (transcriptome profiling) revealed 728 DEGs, with 509 genes differentially upregulated and 219 downregulated in Diff. versus Undiff. cells (**Figure 2A**, left panel). Top 10 DEGs (Upregulated: *DPYSL3, CRABP2, RET, CDH11, BCL2, PRSS12*; Downregulated: *DLK1, RPH3A, EPAS1, IGFBP5*) with the lowest adj. p value were shown in the Volcano plot (**Figure 2A**, right panel). All the DEGs are available in **Supplementary Table 1**.

**Figure 2.**
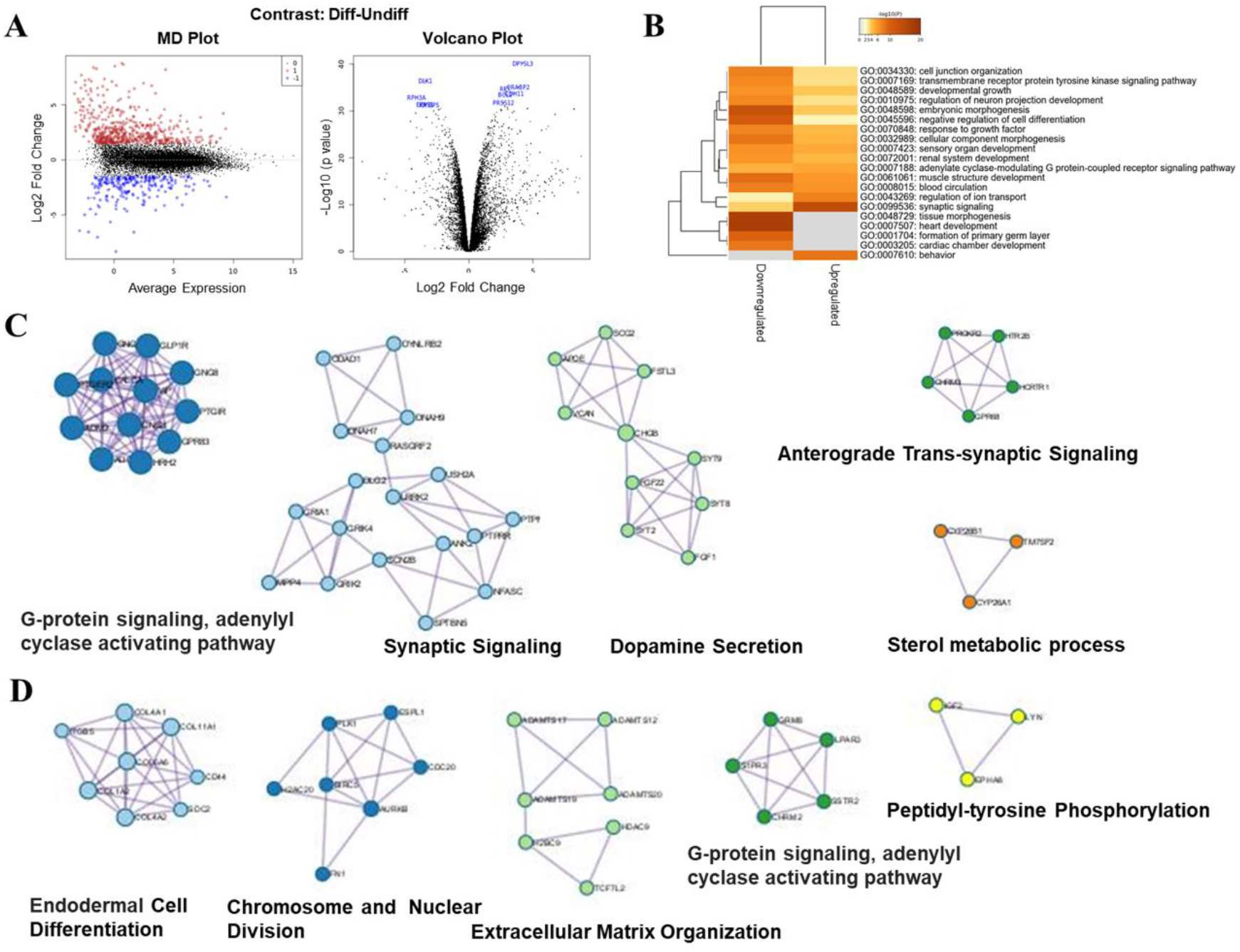
Transcriptomic profiling of neuron-like cells differentiated from SH-SY5Y. **A** The mean difference (MD) plot (left panel) showing the log-fold change and average expression of each gene. Significantly up and down differentially expressed genes (DEGs) are indicated by red and blue dots, respectively. (Cutoff: |Log_2_FC|≥1.5 and adj. P < 0.01). The volcano plot (right panel) displaying the top 10 up and down DEGs with the lowest adj. P value. **B** Heatmap of top 20 enriched Gene Ontology (GO) Biological Process (BP) term across up and down DEGs. **C & D** displaying upregulated (C) and downregulated DEGs (D) Protein-Protein Interaction (PPI) Enrichment Analysis. Each cluster with unique color represents one Molecular Complex Detection (MCODE) components, where the top five p-value terms were retained.

Gene ontology (GO) analysis revealed a strong overlap in enriched GO biological process terms between upregulated and downregulated DEGs, which were significantly involved in synaptic signaling, morphogenesis and developmental pathways (**Figure 2B, Supplementary figure 1A**); GO Cellular Component analysis showed that both upregulated- and downregulated-DEGs were distinctly localized in the neuron-specific structure: synapse, dendrite and axon; The top GO Molecular Function term across all DEGs was calcium ion binding, which functions as a universal second messenger upon which multiple neuronal activities and functions rely on (**Supplementary Figure 1B and C**). These phenomena were in line with expected changes during neuronal maturation and confirmed the outcome of SH-SY5Y neuronal differentiation. To seek the potential interactions of the corresponding gene products, Protein-Protein Interaction (PPI) network analysis was also performed and the five most significant networks for each group were shown in **Figure 2C and D**. Similar to the DEGs, the enrichment of predicted up-regulated proteins were still related to synaptic signaling (Cluster 1 and 4); meanwhile that down-regulated proteins also showed enrichment of GO terms related to cell differentiation (Cluster 1) and cell cycle (Cluster 2). Despite the partial overlap with GO enrichment analysis of DEGs, predicted PPI network clustering exhibited a distinct pattern from DEGs clustering, such as dopamine secretion, further complementing the potential neuron type and function during this process.

### Transcriptome-wide mRNA Stability Profiling of Differentiated SH-SY5Y Cells

Although transcriptomic profiling could reveal the basis of the main morphological and functional alternations during the neuronal differentiation, it still cannot explain some characteristics of these neural-like cells, such as sensitivity to stress. Messenger RNA stability, as a critical element of post-transcription regulation, may be an important mechanism to the complex pattern of neuronal responses to adverse stimuli^17^.

Transcriptome-wide mRNA stabilities were explored by carrying out RNA-seq on Diff. and Undiff. cells treated with ActD for 8 hours. 1619 genes with higher stability (More Stable Genes, MSGs) and 2634 genes with lower stability (Less Stable Genes, LSGs) were identified in the Undiff. group, meanwhile, 1307 MSGs and 2332 LSGs were screened out in the Diff. Group (**Figure 3A**, left panel). Top 10 MSGs and LSGs in Diff. and Undiff. groups were displayed in Volcano plot respectively (**Figure 3A**, right panel). All the MGSs and LSGs are available in **Supplementary Table 2**.

**Figure 3.**
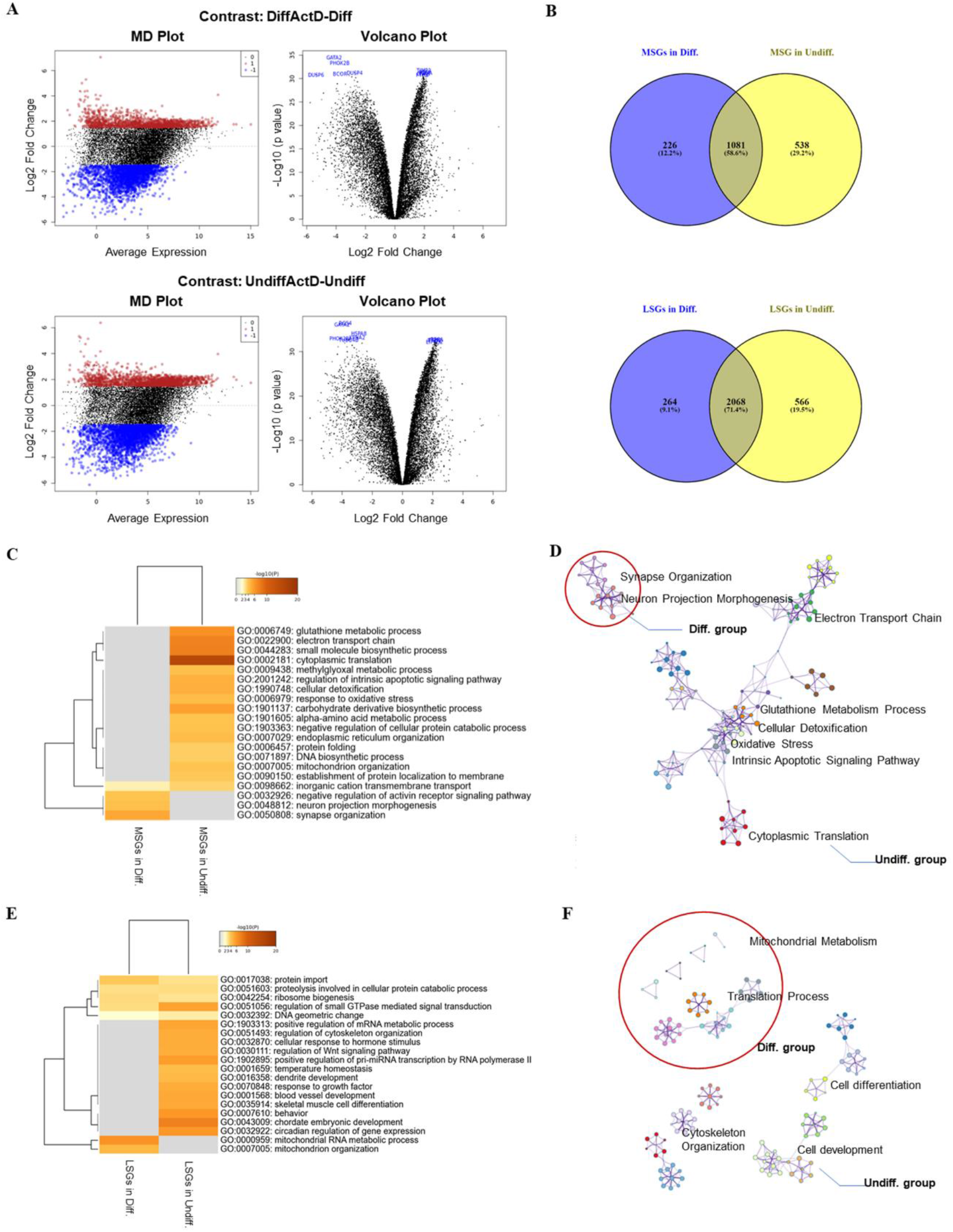
Genome-wide mRNA stability profiling of differentiated and undifferentiated SH-SY5Y cells. **A** The MD plot (left panel) showing the log_2_ FC and average expression of each gene in Diff. (upper panel) and Undiff. group (lower panel). Genes with increased stability (MSGs) and decreased stability (LSGs) are represented by red and blue dots, respectively. (Cutoff: |Log_2_FC|≥1.5 and adj. P < 0.01). The volcano plot (right panel) displaying the top 10 MSGs and LSGs in Diff. (upper panel) and Undiff. group (lower panel) with the lowest adj. P value. **B** Venn Diagram depicting the overlap MSGs (upper panel) and LSGs (lower panel) between Diff. and Undiff. group. **C** Heatmap (colored by p values) and **D** Network (colored by cluster ID) of enriched BP terms of MSGs in Diff. vs Undiff. **E** Heatmap (colored by p values) and **F** Network (colored by cluster ID) of enriched BP terms of LSGs in Diff. vs Undiff. The red circle in D and F refers to clusters enriched in Diff. cells.

Although hundreds of genes exhibited significantly changed stability in both Diff. and Undiff. groups, Venn Diagram displayed the enormous percentage of overlapping MSGs (58.6%) and LSGs (71.4%) between Diff. and Undiff., which could be recognized as gene sets maintaining fundamental cell activities and metabolisms (**Figure 3B**)^18^. Herein, to clearly elucidate the divergence of mRNA stability between Diff. and Undiff., only MSGs and LSGs unique to Diff. and Undiff. groups were employed in the subsequent GO analysis. MSGs in Diff. and Undiff. exhibited an extremely different clustering pattern. In Diff. group, MSGs were still enriched in the pathways related to the morphological changes, such as synapse organization and neuron projection morphogenesis, whereas top GO pathways of MSGs such as cytoplasmic translation, electron transport chain, glutathione metabolic process, cellular detoxification, response to oxidative stress and intrinsic apoptotic signaling pathway were exclusively clustered in Undiff. group. This result gave a clue to the potential molecular mechanism of vulnerability to stress of these neural-like cells (**Figure 3C and D**). Conversely, LSGs in Diff. cells were enriched for protein metabolism and synthesis and mitochondria activity, and top clustered pathways in LSGs of Undiff. cells were cell development and differentiation which accurately represent the phenotypical alterations during the neuronal differentiation (**Figure 3E and F**). Detailed pathway analysis results are available in **Supplementary Figure 2**.

Comparing DEGs with MSGs or LSGs revealed little to no overlap between these datasets (**Supplementary Figure 3, Supplementary Table 3**). This disagreement signifies that mRNA stability changes are independent from transcriptomic changes during neuronal differentiation and represent a unique layer of regulating neuronal function. This is also apparent when examining the enriched GO terms in both datasets (See above).

### Alternative Splicing and RNA binding proteins of Differentiated SH-SY5Y Cells

Alternative splicing (AS), as a key component of post-transcription regulation, enhances proteome diversity by leading to the generation of different protein isoforms, regulating mRNA trafficking, and affecting mRNA stability^19,20^. To decipher the genome-wide splice variations and their potential regulatory functions during neuronal differentiation, differential exon usage (DEU) analyzed by DEXseq and LSV analysis by rMATs were thoroughly explored in differentiated and undifferentiated SH-SY5Y cells.

#### Analysis of Differential exon usage (DEU) by DEXseq

1029 upregulated exons in 766 genes and 2261 downregulated exons in 1535 genes were observed on DEU analysis (**Figure 4A)**. All the DEUs are available in **Supplementary Table 4**. We then performed GO analysis to further investigate the functional enriched pathway clusters of upregulated and downregulated DEUs. The top clusters DEUs enriched in were cellular component morphogenesis, cell junction organization and actin filament-based process (**Figure 4B and C)**, implicating that they all tend to cluster in the pathways related to morphological alterations of neuronal differentiation. Moreover, some downregulated DEUs were distinctly clustered in the mRNA metabolism, peptide biosynthetic process and ribonucleoprotein complex biogenesis, suggesting that the potential regulatory effect of differential exon usage on the mRNA stability, translation, and other epigenetic phenomena may also play a vital role in maintaining the phenotype and function of neuron (**Figure 4B and C**). However, when further comparing DEUs datasets to DEGs and DSGs datasets, the amount of overlap between DEUs and DEGs was much lower than that between DEUs and DSGs (**Supplementary Figure 4A-C**), which further supports the notion that DEUs together with AS and alternative mRNA isoforms are a major factor in impacting the differential mRNA stability after neuronal maturation observed above. Furthermore, GO enrichment analysis of the overlapping genes between DEUs and MSGs/LSGs indicates that many of the top enriched GO terms related to translation process, such as cytoplasmic translation and ribosome assembly in DEU-MSGs pair, and ribonucleoprotein complex biogenesis and protein import in DEU-LSGs pair (**Supplementary Figure 4D and E**), emphasizing that translational processes may be the crucial target of epigenetic/epi-transcriptomic regulators during neuron differentiation.

**Figure 4.**
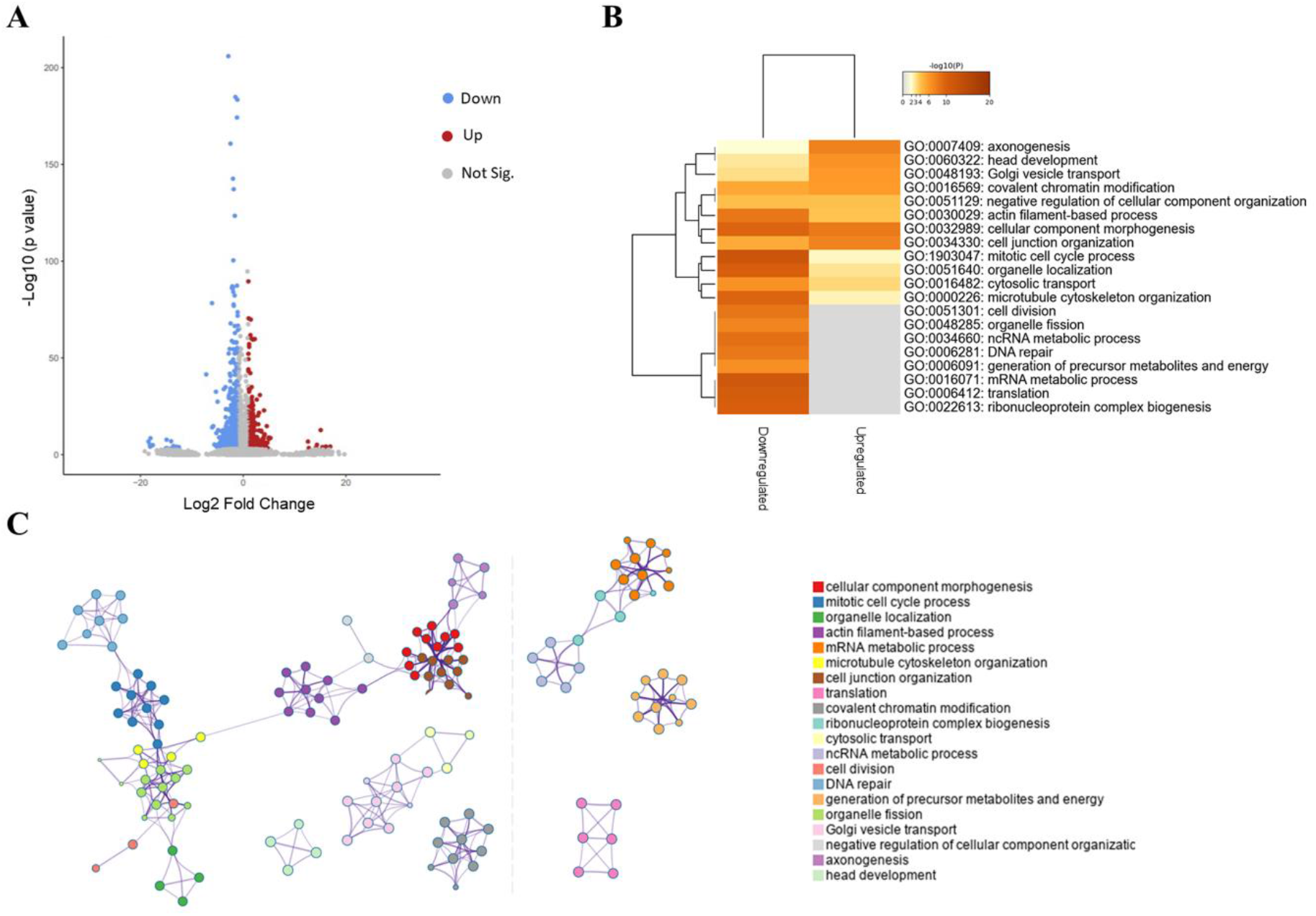
Differential exon usage (DEU) by DEXseq of differentiated and undifferentiated SH-SY5Y cells. **A** Volcano plot showing differential exon usage between Diff. and Undiff. group. Red dots for upregulated DEUs, and blue dots for downregulated DEUs. **B** Heatmap and **C** Network (colored by cluster ID) of enriched BP terms of DEUs in Diff. vs Undiff. group.

#### Analysis of local splicing events (LSVs) and RNA-binding proteins (RBPs) motif enrichment

Differential exon usage can give a good idea about mRNA isoforms expression; however, it fails to grasp the complexity of the alternative splicing (AS) program. To that end, we employed the rMATs pipeline to analyze local splice variants (LSVs) and evaluate the differential enrichment of different AS programs^21^. rMATs identified a total of 5594 differential alternative splicing events in our dataset. SE (3336 events, 60% of all LSV events) was the most prevalent AS event compared to other 4 types of AS (MXE: 889, 16%; A3SS: 538, 9%; A5SS: 436, 8%; RI: 395, 7%) (**Figure 5A-F)**. All the LSVs are available in **Supplementary Table 5**.

**Figure 5.**
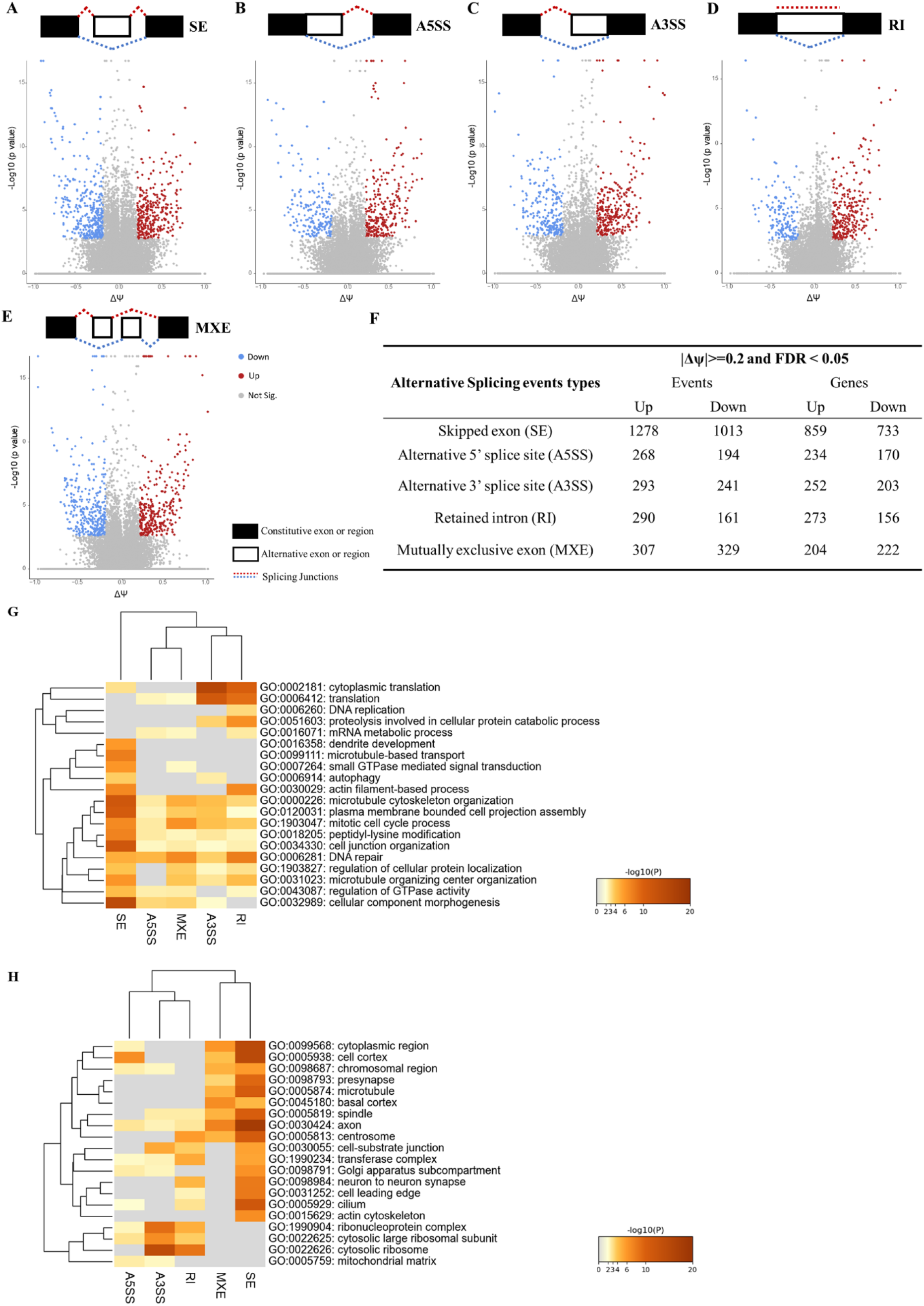
Alternative Splicing (AS) events by rMATs in differentiated and undifferentiated SH-SY5Y cells. **A-E** Volcano plots showing differential skipping exons (SE), alternative 5’ splice site (A5SS), alternative 3’ splice site (A3SS), retained intron (RI) and mutually exclusive exon (MXE) with the corresponding schematic diagram of each AS events. Red dots for upregulated and blue dots for downregulated AS events. **F** Summary of the number of AS events and related genes. Heatmap (colored by p values) of top 20 BP term (**G**) and cellular component term (**H**) of each differential AS events in Diff. vs Undiff.

GO pathway analysis demonstrated that SE events, and to a lesser extent other LSV types, were functionally enriched for morphological related pathway, such as cellular component morphogenesis, plasma membrane bounded cell projection assembly, microtubule cytoskeleton organization and cell junction organization, in accordance with DEGs and DEUs enriched biological pathways. It is noted that the other four AS events, A5SS, MXE, A3SS and RI, exclusively clustered in mRNA metabolism and translation (**Figure 5G**). The distinct cluster pattern of the other 4 AS events suggested the potential effect of alternative splicing on the translation during the neuronal differentiation process. Furthermore, GO cellular component pathway analysis reinforced this phenomenon: differential SE events were specially enriched for genes related to the axon, where major morphological changes frequently take place during the neuronal differentiation. On the contrary, genes regulated by A5SS, MXE, A3SS and RI were more linked to the ribosome and mitochondria (**Figure 5H**). This pattern of dichotomous enrichment, and the concordance of SE with DEUs (which also represent SE events) and DEGs indicate that different aspects of neuronal functional specialization occur during neuronal differentiation and maturation. It also signifies the importance of AS reprogramming in this process as well as in maintaining neuronal functions. Additionally, the comparison of each AS events to DEGs and DSGs datasets showed consistency with the observation of DEUs: each AS events-DSGs pairs exhibited larger overlap than AS events-DEGs pairs (**Supplementary Figure 5 and 6**), signifying the regulatory effect of AS on mRNA stabilization during the neuronal differentiation.

**Figure 6.**
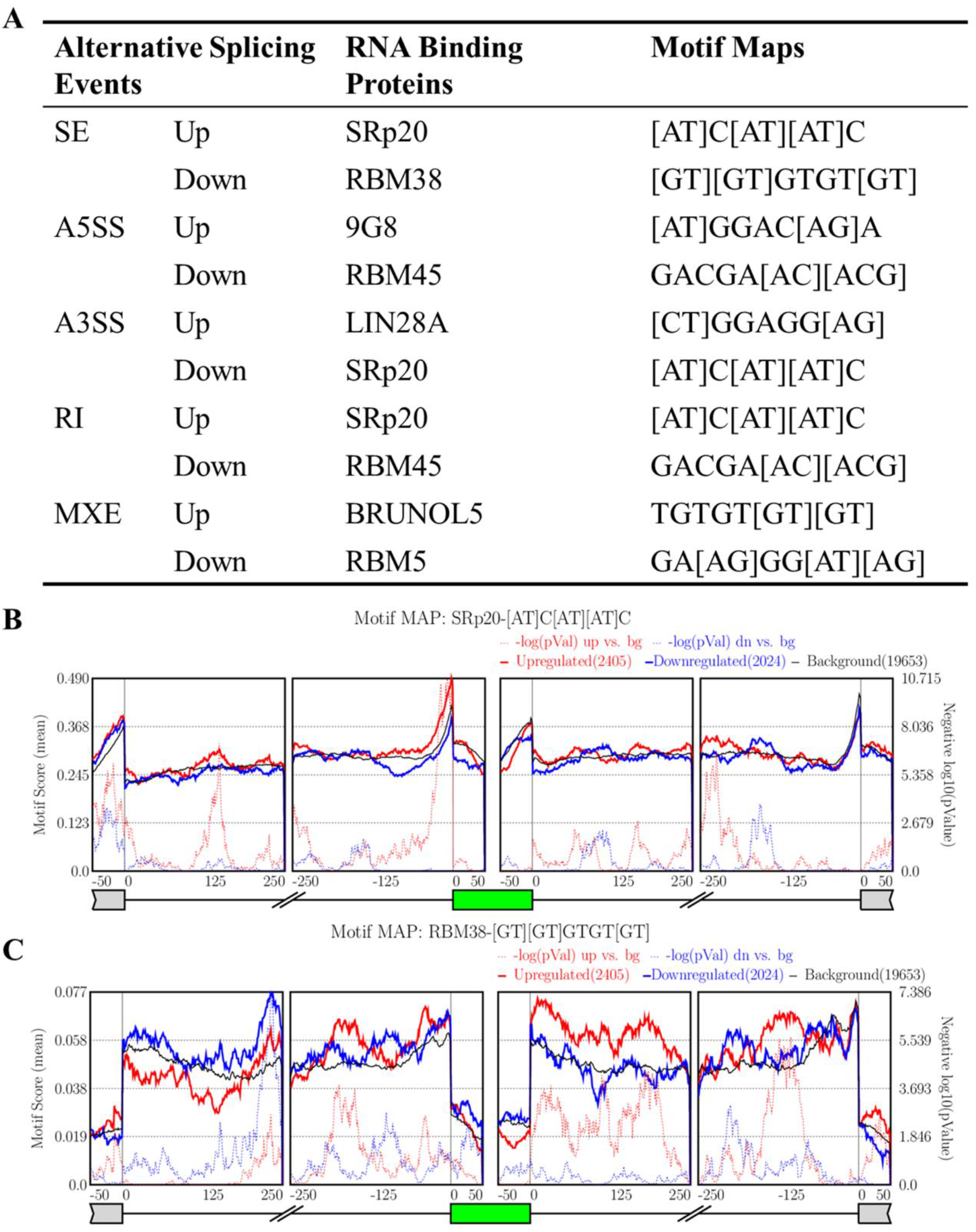
Prediction of RNA binding proteins (RBPs) within each alternative splicing event by rMAPS2 in differentiated vs undifferentiated SH-SY5Y cells. **A** Summary of the most significantly enriched RPBs motif maps for each up and down alternative splicing events. **B** RNA motif maps showing enrichment of SRp20 near alternatively spliced exons. **C** RNA motif maps showing enrichment of RBM38 near alternatively spliced exons. The dotted lines indicate the significance of enrichment versus background. The solid lines indicate the motif score of upregulated and downregulated genes compared to background genes.

RNA-binding proteins (RBPs) bind to pre-mRNAs and mRNAs and thereby tightly control AS or mRNA translation and stability, in a cell-type-specific manner, especially in neurons^22,23^. To identify the explicit RPBs enriched for each AS events and evaluate their possible biological effect in differentiated SH-SY5Y cells, motif enrichment analysis using rMAPS2 was performed with differential AS data generated from rMATS. The most significantly enriched RBPs for each up- and down-regulated AS events are displayed in **Figure 6A**. Interestingly, SRp20 appeared to be the most common RBPs among the list of top enriched RBPs, which was also shared by upregulated exon with SE and RI and downregulated exon with A3SS, suggesting its essential regulatory role during the neuronal differentiation. Due to the fact that SE event is the most prevalent AS events in the human genome, we focuses on top enriched RBPs (SRp20 and RBM38) for SE in this study^24^. SRp20 binding motifs were enriched in the upstream intron and the downstream intron of the upregulated exons (solid and dotted red line show peaks) and in the downstream intron of the downregulated exons (solid and dotted blue lines show peaks) (**Figure 6B**) This result is consistent with the published mRNA binding properties of SRp20^25^. Furthermore, a strong enrichment for RBM38 motif were observed within the downstream intron regions when the inclusion of the target exon is promoted (solid and dotted red line show peaks), conversely, it was enriched in the upstream flanking intron regions and target exons when the inclusion of the target exon is suppressed (solid and dotted blue line show peaks), indicating the potential regulatory mechanism of RBM38 on SE in the context of neuronal differentiation (**Figure 6C**). The rest RNA motif maps of most enriched RBPs for A5SS, A3SS, RI and MXE could be available in **Supplementary Figure 7**.

**Figure 7.**
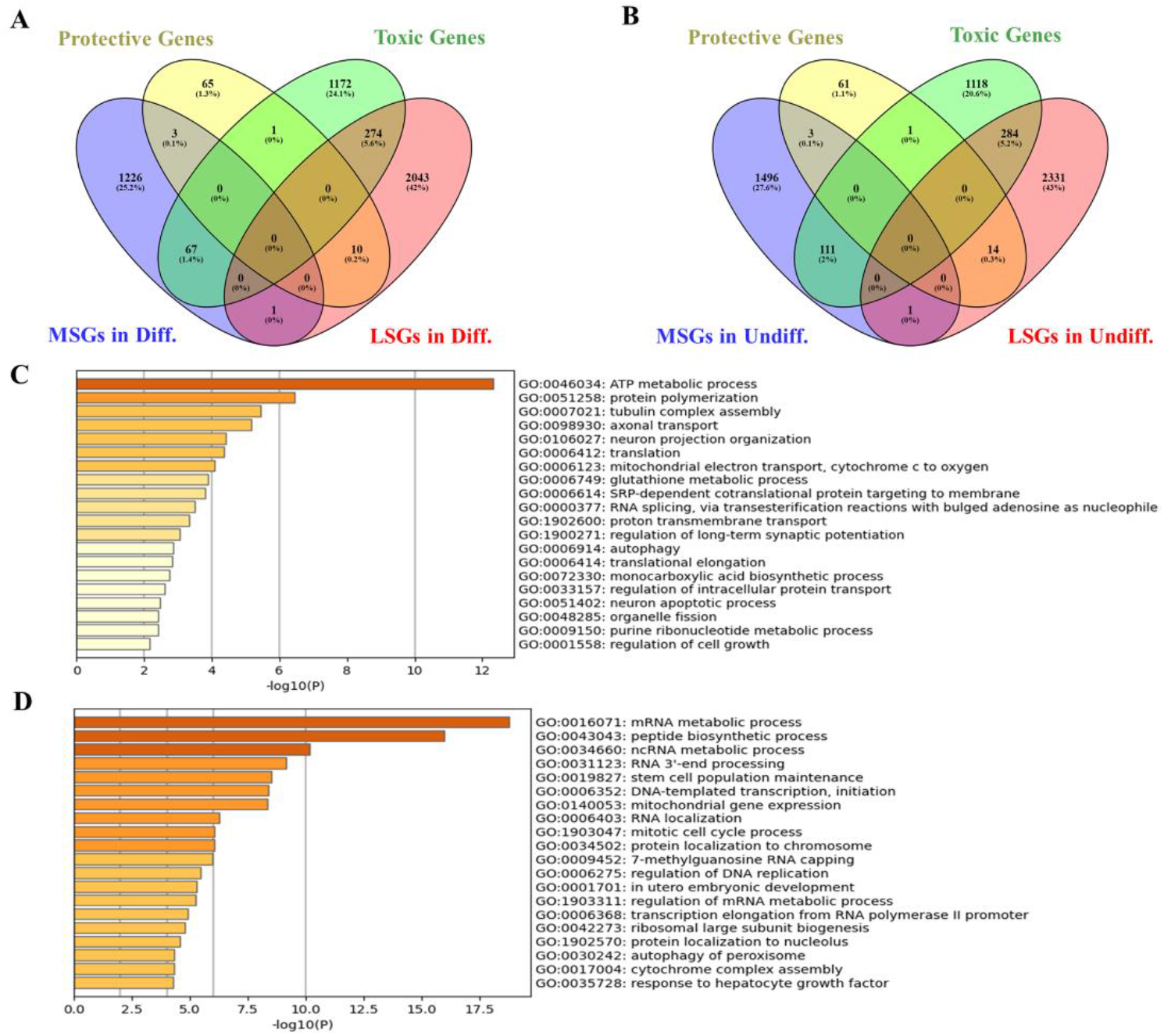
Integrative analysis between mRNA stability profiling of both differentiated and undifferentiated SH-SY5Y cells and genomic-wide CRISPRi screens on iPSC-derived neurons under the chronic oxidative stress. Venn diagrams displaying the ratio of overlap between genes with altered stability in Diff. (**A**) and Undiff. (**B**) SH-SY5Y cells and survival genes of iPSC-derived neurons. Bar graph showing enriched BP terms of Toxic-MSGs (**C**) and Toxic-LSGs (**D**), colored by p-values.

### Integrative analysis between CRISPRi Screens of iPSC-derived neurons and transcriptomic analysis of SH-SY5Y cells

A recent study conducted genome-wide CRISPR screens on iPSC-derived neurons to identify genes linked to neuronal fitness and neuronal survival, providing a treasure trove of data to understand neurons’ molecular and genetic landscape^15^. CRISPR inference (CRISPRi) and CRISPR activation (CRISPRa) were performed on human neuron under chronic mild oxidative stress and control conditions and identified the top hit genes that are functionally linked to oxidative stress sensitivity and cell death in neurons, suggesting the selective vulnerability of neuron. To further verify the potential link between mRNA stability and the selective sensitivity to stress of neuron, we performed two comparisons between transcriptomics and mRNA stability changes of differentiated SH-SY5Y cells and survival genes dataset of iPSC-derived neurons. First, DEGs of SH-SY5Y cells were hard to match the survival genes from CRISPRi or CRISPRa screens (**Supplementary Figure 8, Supplementary Table 6**), suggesting the differential gene expression of neuron may not be the reason of neuron specific vulnerability.

Next, we compared the mRNA stability profiling datasets and CRISPRi screen under the mild oxidative stress. The comparison between DSGs datasets and CRISPRi showed much stronger overlap than that between DEGs and CRISPRi (**Figure 7A and B**). Next, we performed GO pathways analysis on the overlapping datasets of Toxic-MSGs (Toxic genes that promote oxidative stress/cell death) and Toxic-LSGs, which showed higher overlapping ratio compared to other groups. Interestingly, the most significantly enriched GO biological processes terms of Toxic Genes-MSGs was ATP metabolic process, reinforcing the link between altered mRNA stability and mitochondria dysfunction and subsequently redox imbalance in neuron^26^ (**Figure 7C**). It was intriguing to find that genes of Toxic Genes-LSGs dataset were significantly enriched in mRNA metabolic process and peptide biosynthetic process, indicating the possible regulatory role of mRNA decay on the translation in the context of neuron lineage progression (**Figure 7D**).

## Discussion

To decipher the explicit molecular mechanism during the neuronal differentiation and further interpret the neuron behavior, we performed deep RNA-seq on differentiated and undifferentiated SH-SY5Y cells. Our data revealed that the acquisition and maintenance of neuron-specific structures and functions may be co-regulated by an intricate network consisting of mRNA transcription, stability, splicing, and splicing factors/RBPs. Interestingly, dynamic changes in mRNA stability appears to be the only regulatory factor contributing to the selective vulnerability of mature neurons to oxidative stress. The profiling of AS events and the correlated RBPs provides a rich resource for complementing our understanding of neuron differentiation and maturation.

### Transcriptomic changes during the neuronal differentiation

In this study, we showed that SH-SY5Y cells treated with RA and BDNF could switch from an undifferentiated state towards a mature neuron-like phenotype, mimicking the process of neuronal differentiation. Transcriptomic profiling on differentiated SH-SY5Y cells disclosed that abundant DEGs are closely related to neuronal morphogenesis and function (i.e.: synaptic signaling), which is in line with previous studies on differentiated SH-SY5Y cells^12,13^. Similar changes of gene expression pattern were also observed in human iPSC-derived neurons^27,28^ and human ESC-derived neurons^29^, confirming the validity of SH-SY5Y as neuronal differentiation model. Moreover, our PPI networks identified that predicted up-regulated proteins in differentiated SH-SY5Y cells still related to synaptic signaling and down-regulated proteins involved in cell differentiation and proliferation, consistent with previous global proteomic analysis of differentiated SH-SY5Y cells^30^. To be noted, DEGs and their corresponding predicted proteins shared similar GO enrichment patterns, indicating that a coordinated transcriptomic and proteomic reprogramming of SH-SY5Y cells into neuron-like cells. However, the apparent difference between the protein clusters and DEGs clusters was also observed, reflecting possible regulatory mechanisms on the protein level at play^31^. Additionally, epigenetic regulation might also be an explanation to this phenomenon^32,33^. Taken together, although our bioinformatic analysis required further experimental verification, these transcriptomic data in differentiated SH-SY5Y cells still provide a rich resource to advance our understanding of the molecular mechanism involved in neuronal differentiation and maturation.

### mRNA stability alternation during the neuronal differentiation

Steady-state RNA levels during the neuronal differentiation are maintained by a balance of RNA transcription and decay^34^. By inhibiting transcription using ActD, measuring RNA abundance can reflect mRNA stability and indirectly represent the translation efficiency^35,36^. In this study, we conducted the first global analysis of mRNA stability on undifferentiated and differentiated SH-SY5Y cells by deep RNA-seq, unraveling its essential role in controlling neuronal differentiation. Although DSGs dataset is significantly different from DEGs dataset, it is still closely involved in known neural-specific pathways, indicating that the altered stability may support appropriate levels of neuron-related protein production without the requirement of high/low rates of transcription. In differentiated SH-SY5Y cells, MSGs functionally belong to the GO categories of neuron projection morphogenesis and synapse organization, in accordance with the phenotype of mature neuron; LSGs predominantly cluster in ribosome biogenesis and mRNA metabolism, both of which is tightly linked to cell growth, proliferation, and differentiation^37,38^. This may relate to the transition from proliferation to differentiation and maturation that occurs in SH-SY5Y cells. This divergent pattern of mRNA stability has also been observed in the nervous system of Drosophila embryo^8^. Destabilization of mRNAs related to the transcription and translation enabled cells to rapidly respond to differentiation inducers and produced dynamic molecules to initiate differentiation process; Selective stabilization of neuron-specific mRNAs could be helpful to initiate the process of neuronal specification and maturation^39^. This observation provides a new angle to interpret the neuronal differentiation, which could be further investigated on neuronal lineage progression of stem cell.

### Altered mRNA stability may be the critical mechanism of selective vulnerability to oxidative stress of neuron

Besides utilized as neuronal differentiation model, SH-SY5Y cells were widely employed as *in-vitro* model for neurodegenerative diseases, especially Parkinson’s disease, due to its susceptibility to mitochondrial dysfunction and high vulnerability to oxidative stress after differentiation^40^. However, existing studies on the vulnerability of differentiated SH-SY5Y cells focused on the phenotypic verification or mechanism exploration in low throughput approaches^14,40^ leading to the limited explanation of this heightened sensitivity of oxidative stress. Here, our mRNA stability profiling collectively unraveled exclusive stabilization of genes related to redox homeostasis in the undifferentiated cells and destabilization of genes involved in mitochondria metabolism and function under the differentiated state. Most interestingly, our differentially stabilized gene sets partially overlap with survival genes controlling neuronal response to chronic oxidative stress uncovered by CRISPRi screen on iPSC-derived neurons^15^. This is the first-hand data to disclosed that mRNA stability dynamics could serve as an essential mechanism of selective vulnerability of neuron.

Although the causal link between DNA methylation and histone modification (two main epigenetic regulation) and neurodegenerative disorders are well discussed, how RNA based epigenetic regulation contributes to vulnerability to adverse stimuli of neuron is an emerging question^32^. mRNA stability can be impacted via many mechanisms such as AS, mRNA translation rates, mRNA Modifications, and others^17,41-43^. In physiological states, RNA homeostasis is the outcome of the intricate balance between stability promoting factors (5-capping/3-polyadenylation modification) and decay factors (RNA-degrading enzymes), both of which are under the control of RBPs^44^. This balance is of particular importance in neurons, which are among the most metabolically active and morphologically complex cells, and any disruptions in this balance could exert dramatic consequences for neuron viability. For example, neuron specific RBPs HuB, HuC and HuD could enhance RNA stability by upregulating alternative polyadenylation and loss of them could sensitize neuron to oxidative stress^45,46^. On the other hand, when exposed to stress, stress granules (SGs) and processing bodies are two competing factors to the sequester and stabilize mRNAs altering their translation or degradation rates^44^. The formation of SGs occurs immediately upon exposed to stress, consequently sequestering and stabilizing some essential mRNAs in SGs suppressing their translation^47,48^. However, neurons possess a greater diversity in SGs composition than non-neuronal cell types, frequently exhibit aberrant SGs especially in the disease state, leading to destabilization of survival genes and cause a series of downstream toxicity ^49^. Although our result regards mRNA stability as a novel insight into understanding the selective vulnerability of neuron, the multiple regulators of mRNA stability need to be further explored and more complete investigations on RNA degradation under both normal and stress conditions will be necessary.

### The role of alternative splicing during the neuronal differentiation

Alternative splicing occurs at high frequency in brain tissue and may contribute to various biological process in the nervous system^50^. To understand in depth the likely biological consequences of AS patterns in the context of neuronal differentiation, mRNA isoforms and specific alternative splicing events on differentiated and undifferentiated SH-SY5Y cells were globally investigated through deep RNA-seq, recognizing abundant exons and splicing events occurring in the neuronal differentiation. Gene ontology analysis revealed some clues on what the function of these differential AS event during neuronal differentiation might be. Among transcripts with differential exon usage, cellular component morphogenesis and mitotic cell cycle process were the most prominent in upregulated and downregulated gene sets respectively, suggesting differential exon usage might be involved in cell cycle withdrawal and differentiation initiation, and the induction of morphological alterations including neurite outgrowth and/or synapse formation. The top emergence of the same pathway of skipped exons (SE) events appeared to be consistent with this finding^50,51^. These results consistently verify the indispensable role of AS in the neuronal lineage progression.

It was intriguing to discover the dimorphic enrichment pattern between SE and the other AS events including A5SS, A3SS, RI and MXE. The latter four AS events were closely involved in the translation regulation and localized in the ribosome/ribonucleoprotein complex, suggesting that A5SS A3SS, RI and MXE may have the dominance in regulating epigenetic and translational processes these neuron-like cells. Taking RI as example, the existing evidence showed that RI in the 5’ untranslated region (UTR) has the potential to regulate translation initiation via conferring sensitivity to 4E-BP-mediated translational suppression during the early phases of embryonic development^52^. How AS would regulate translation in a LSV type specific/cell type specific manner still require further investigation, our study collectively presented a characterization of different alternative splicing events during the neuronal differentiation.

It is well recognized that AS regulates gene expression and is at the base of phenotypic alternation. However, the AS genes in our study did not overlap with gene expression profiling, indicating AS may have a limited effect on changing RNA abundance in the context of this work. This phenomenon brings about another question: how does AS participate in the transformation from undifferentiated to differentiated without interfering with transcriptome? One possible explanation is the direct effect of AS on translational efficiency, shown in the aforementioned discussion. Moreover, based on the high similarity between our AS profiling and stability profiling, AS may have a priority effect on mRNA stability via mRNA-RBPs interactions or by yielding different mRNA isoforms that differ drastically in their half-lives and translational output. Previous study shows that the human 5-HT1A mRNA 3’-UTR is alternatively spliced to generate an ultra-stable isoforms which is efficiently translated in a neural-specific manner^53^, providing the evidence of the regulatory role of AS on RNA stability in neuron. These results presented here may be a useful clue to explore which the dominant mechanism that AS will choose to regulate different neuronal phenotypes or activities.

The aforementioned mechanisms such as mRNA splicing, and stability are tightly regulated by RBPs. particularly in neurons. Therefore, we globally analyzed enriched RBPs motif maps for each AS events and screened out the most significant changed RBPs SRp20, which may emerge as a crucial regulator of different AS events to regulate neuronal differentiation process. SRp20, as the smallest member of Ser/Arg-rich splicing factor family, could promote the exclusion/inclusion of exon to realize its regulation on cellular proliferation and/or maturation. Dysregulated SRp20 expression has been implicated in the pathophysiology of neurodegenerative disorder Alzheimer’s disease by splicing TRKB^54-57^. Our data implied that beyond the SE splicing, SRp20 may exert on A5SS and RI to fine tune the process of neuronal differentiation. However, its potential target gene locus and its corresponding biological function need more effort to verify. Moreover, SRp20, by interacting with the cellular RNA-binding protein, PCBP2, becomes a necessary mechanism for translation initiation of viral RNAs^58^. Although the explicit effect of SRp20 in the neuronal differentiation remains to be further explored, our finding firstly points out that SRp20, emerging as a new neuron-specific splicing factor, may have a universal effect on different AS events.

### Limitations

There are some limitations of this study that need to be acknowledged. The main findings are concluded based on the bioinformatic analysis, which usually needs further experimental verification. However, due to the identification of various processes and the huge dataset generation during the course of this work, exploring all of them will make this study exhaustive and beyond the scope of a single article. Future efforts will be imperative to analyze the various processes described herein and validate their potential roles in neuronal differentiation and functioning.

## Conclusion

In summary, our quantitative profiling of transcriptome and mRNA stability and alternative splicing demonstrated that the sequential consequence of neuronal differentiation involves transcriptional, post-transcriptional and epigenetic re-programming. The heightened sensitivity to stress of mature SH-SY5Y cells after differentiation may be tightly orchestrated by epigenetic factors, which is a novel insight into interpreting selective vulnerability to adverse stress of neuron. With the integrative analysis of this regulatory network, our data may broaden our understanding on the molecular basis of neuronal differentiation and behavior and could serve as a paradigm to comprehensively analyze RNA metabolism in neurons.

## Methods Cell culture

Human SH-SY5Y neuroblastoma cells obtained from ATCC (Cat# CRL-2266) were cultured in Eagle’s Minimum Essential Medium (EMEM; ATCC, Cat# 30-2003) and F-12 Nutrient Mixture (HAM; Gibco, Cat# 11765054) 1:1 containing 10% heat-inactivated Fetal Bovine Serum (FBS; Corning, Cat# 27419002), at 37°Cand 5% CO2. No antibiotics were added to the growth media.

### Differentiation protocol

The presented neuronal differentiation protocol for SH-SY5Y cells was modified from Enicinas et al ^11^. The differentiation protocol consisted of two stages; a 4-day pre-differentiation step in EMEM/F-12 supplemented with 10% FBS and 10uM RA (Sigma-Aldrich, Cat# R2625), and a subsequent 6-day differentiation step in serum-free medium containing 50 ng/ml human brain derived neurotrophic factor (BDNF) (Sigma-Aldrich, Cat# B3795). Cells were seeded at an initial density of 2 × 10^4^ cells/cm^2^ in 24 well plate coated with Type I collagen (Corning, Cat# 3524). Media were routinely changed every 2-3 days. Differentiation efficiency was evaluated by observing morphological alterations under phase contrast microscope (Leica DMi1) and immunofluorescent staining for neuronal markers.

### Immunofluorescent staining

Cells seeded on 8 well glass slide (Millipore, Cat# PEZGS0816) coated with poly-L-lysine (Sigma-Aldrich, Cat# P6282) were fixed with 2% paraformaldehyde (PFA, Fujifilm Wako, Cat# 162-16065) in phosphate buffered saline (PBS) for 30 min at 4°C. Cells were permeabilizated with 0.1% Triton X-100 (Nacalai Tesque, Cat# 35501-15) for 10 min and then blocked by 2% bovine serum albumin (BSA) (Nacalai Tesque, Cat# 01860-07) for 1h at 4°C. Cells were then incubated overnight with primary antibodies Nestin (Biolegend, 839801, 1:200), MAP-2 (Santa Cruz, Sc-5359, 1:50), β-tubulin-III (Cell signaling, 2128, 1:200), or Synaptophysin (Abcam, Ab14692, 1:100) diluted in PBS containing 1% BSA and 0.1% Triton-X-100 at 4 °C. Anti-rabbit and anti-goat secondary antibodies conjugated with Alexa-Fluor 488 (Invitrogen, 1:2000, A11034) or Alexa-Fluor 555 dyes (Abcam, 1:2000, ab150130) diluted in the same antibody buffer were incubated with the cells for 2h at 4 °C. Cells were washed once in PBS and counterstained with 4′,6-diamidino-2-phenylindole di-hydrochloride (DAPI, Vector, Cat# H-1500). Images were taken using a laser confocal microscope (Olympus IX83) and analyzed with FLUOVIEW FV3000.

### RNA isolation and quality control

Cells were lysed in QIAzol Lysis Reagent (QIAGEN, Cat# 79306) for RNA extraction. RNA was extracted using the miRNeasy Kit (QIAGEN Cat# 217004) with a DNase digestion step following the manufacturer’s instructions. RNA purity and concentration were examined by Nanodrop (Thermo Fisher Scientific; Catalog# ND-ONE-W), and RNA integrity number (RIN) was assessed using RNA 6000 Nano Kit (Agilent, Cat# 5067-1511) on Agilent Bioanalyzer 2100. Samples with RNA integrity number ≥ 9 were used.

### RNA sequencing

RNA-seq libraries were prepared using NEBNext Poly(A) mRNA magnetic isolation module (for mRNA enrichment) and Ultra RNA Library Prep Kit (NEB, Cat# E7760) following the manufacturer’s instruction. Library quality was assessed by Agilent DNA 1000 kit (Agilent, Cat# 5067-1504)) on Agilent Bioanalyzer 2100. Libraries concentration was determined using qPCR using NEBNext library Quant kit for Illumina (NEB, Cat# E7630). Libraries were pooled and sequenced by Macrogen on Ilumina Hiseq X-ten. Sequencing was performed with 150 base-pair pair-end reads (150bp × 2). Samples were sequenced at a target depth of 50 million reads per sample.

### Messenger RNA stability profiling by Actinomycin D assay

Actinomycin D (ActD) (Sigma-Aldrich, Cat# A9415), a transcription inhibitor, is widely used in mRNA stability assays to inhibit the synthesis of new mRNA, allowing the evaluation of mRNA decay by measuring mRNA abundance^59^. Cells were treated with 5ug/ml ActD for 8 h, and then collected for RNA-seq.

### RNA-seq data analysis

#### Differential Expression and mRNA stability Analysis

Raw Fastq files were first trimmed with Trimmomatic^60^ and aligned to the human reference genome hg38 (GRCh38.p13) using the splice aware aligner HISAT2^61^. Around 84.5%-87.5% of read pairs were uniquely mapped to the hg38 genome. Mapped reads BAM file were then counted to gene features by FeatureCounts^62^ with standard settings. Differentially expressed genes (DEGs) and genes with altered RNA stability (defined as |Log_2_ FC ≥ 1.5 and adj. p value < 0.01) were analyzed by Limma^63^ with normalization method TMM. For the mRNA stability analysis, we compared differentiated (Diff) and undifferentiated (Undiff) datasets to their ActD treated counterparts. We hypothesized that if the stability of mRNAs is equal, we shall not see changes. However, the existence of DEGs in this comparison indicates changes in stability of mRNAs in relation to the total RNA pool. Next, we compared the stabilized and destabilized mRNAs between Diff and Undiff groups to get the differentially stabilized genes (DSG).

This analysis was performed using an Online instance of Galaxy^64^.

#### Differential exon usage analysis by DEXseq

The analysis on differential exon usage (DEU) was also performed using Galaxy. First, we counted exons using DEXseq-counts. Then, DEU was analyzed by DEXseq^65^ with FDR < 0.05 and |Log_2_ FC| ≥ 1 as cutoff values.

#### Local splice variants analysis by rMATs and RNA Binding Protein (RBP) Motifs analysis by rMAPS2

Alternative splicing was analyzed with rMATS using the ENSEMBL gene models (Homo Sapiens.GRCh38.p13.gtf)^21^. rMATs can detect 5 different local splice variants (LSVs) skipped exons (SE), mutually exclusive exons (MXE), alternative 3’ and 5’ splice sites (A3SS and A5SS respectively) and retained introns (RI)). We used FDR < 0.05 and Deltapsi (ΔΨ) ≥ 0.2 as cutoff values for significant splice events.

RNA binding proteins (RBPs) motif enrichment from differentially regulated AS events between Diff. and Undiff. group were analyzed by online software rMAPS2 (http://rmaps.cecsrearch.org/)^66^. The output files of five AS events obtained by rMATS were used as input for rMAPS2 and hg38 was selected as reference genome assembly.

### Functional Enrichment Analysis

Functional enrichment analysis was carried out to detect the most enriched Gene Ontology (GO) terms in gene sets obtained by various analysis methods. All the genes with significant changes from each analysis were analyzed by Metascape ^67^.

## Data availability

The raw sequencing data files are available through Sequence Read Archive (SRA) database with accession number PRJNA779467.

## Author contribution

**ZY**: Performed all experiments. Data analysis and interpretation. Wrote the manuscript. **SR**: Conception and study design. Data analysis and interpretation. Critically revised the manuscript. Funding and administration. Study supervision. **TT**: Critically revised the manuscript. **KN**: Critically revised the manuscript. Study supervision.

### Funding

This study was funded by Japan society for promotion of science grants number 20K16323 and 20KK0338 for SR.

## Acknowledgment

The authors report no conflict of interest nor there are any ethical adherences regarding this work.

