## Supplementary figures for "Mature Neurons’ sensitivity to oxidative stress is epigenetically programmed by alternative splicing and mRNA stability"

### Lead author

**Corresponding authors:**

Sherif Rashad:

Kuniyasu Niizuma:

### Supplementary figures

**Supplementary Figure 1** Transcriptomic analysis of differentiated and undifferentiated SH-SY5Y cells by Metascape. **Related to Figure 2**. **A** Network of enriched terms of differentiated SH-SY5Y cells, colored by each cluster. **B** Heatmap of enriched Cellular Component (CC) terms across Up and Down DEGs. **C** Heatmap of enriched Molecular Functions (MF) terms across Up and Down DEGs.

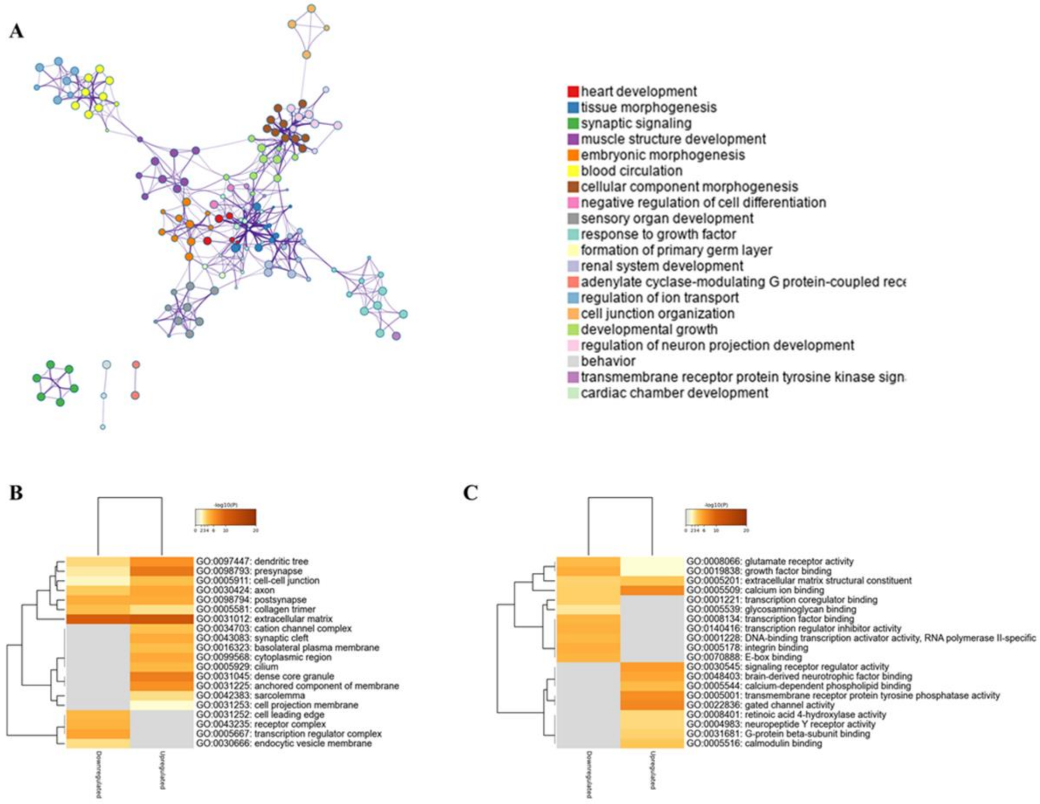

**Supplementary Figure 2** Genome-wide mRNA stability profiling of differentiated and undifferentiated SH-SY5Y. **Related to Figure 3**. **A** Network (colored by p value) of enriched BP terms of MSGs in Diff. vs Undiff. **B** Network (colored by group enrichment level (Diff vs Undiff)) of MSGs enriched BP pathways showing the distinct pattern between Diff. and Undiff. **C** Network (colored by p value) of enriched BP terms of LSGs in Diff. vs Undiff. **D** Network (colored by group enrichment level) of LSGs enriched BP pathways showing the distinct pattern between Diff. and Undiff.

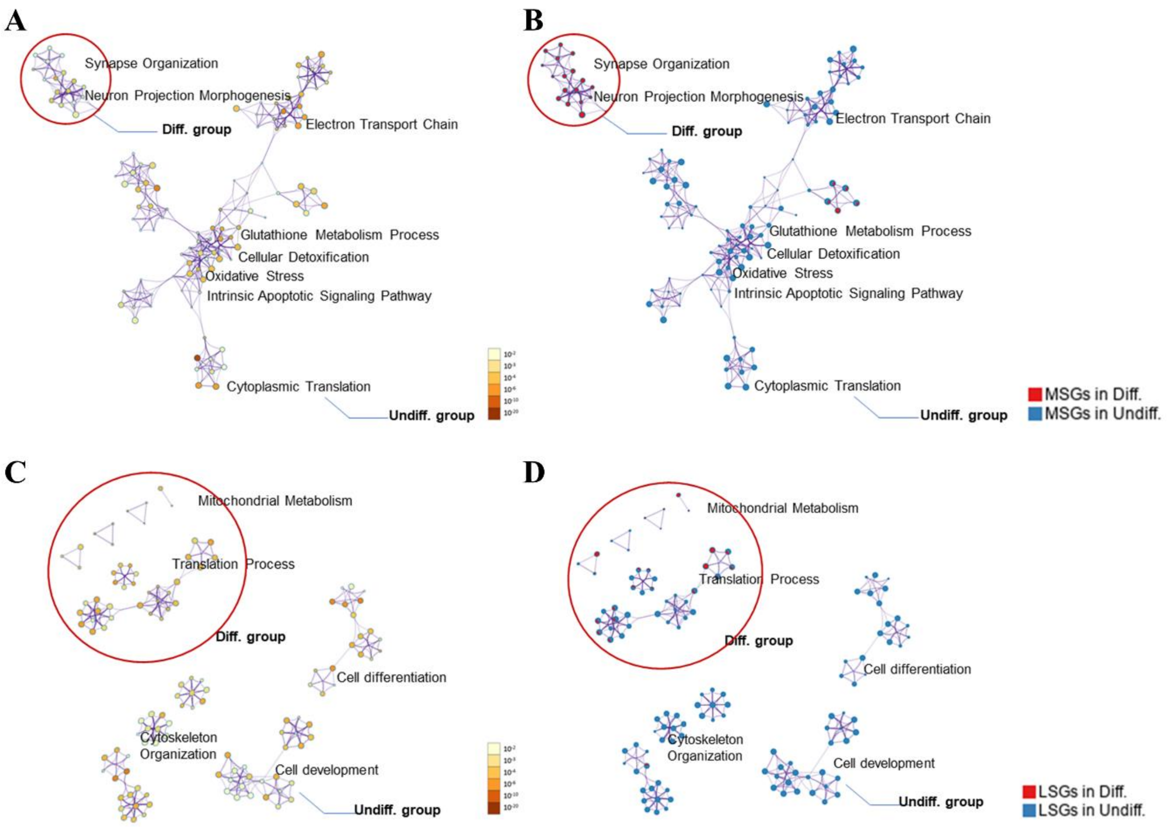

**Supplementary Figure 3** Comparative analysis between transcriptomics and mRNA stability data of differentiated SH-SY5Y cells. **Related to Figures 2 and 3.** Venn diagrams showing the percentage of overlap of DEGs and MSGs/LSGs in Diff. (**A**) and Undiff. (**B**) of differentiated SH-SY5Y.

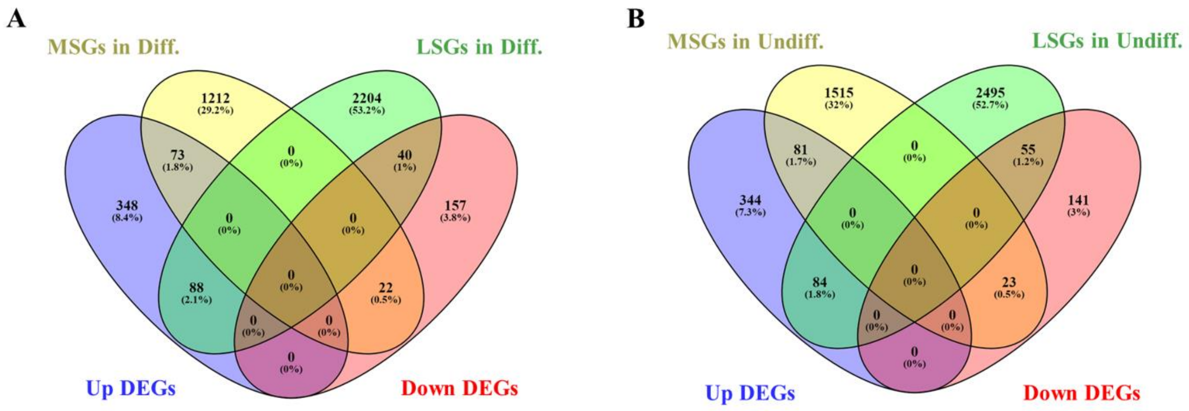

**Supplementary Figure 4** Comparative analysis of differential exon usage to transcriptomics and mRNA stability profiling in SH-SY5Y cells. **Related to Figures 2, 3 and 4**. **A** Venn diagram presenting the overlap between DEGs and DEUs. Venn diagram depicting the overlap between mRNA stability profiling and DEUs in differentiated (**B**) and undifferentiated (**C**) SH-SY5Y cells. Heatmap showing top 20 significantly enriched GO terms of overlapping DEU-MSGs (DEU here refers to all DEUs and upregulated or downregulated) (**D**) and DEU-LSGs (**E**) in Diff and Undiff cells.

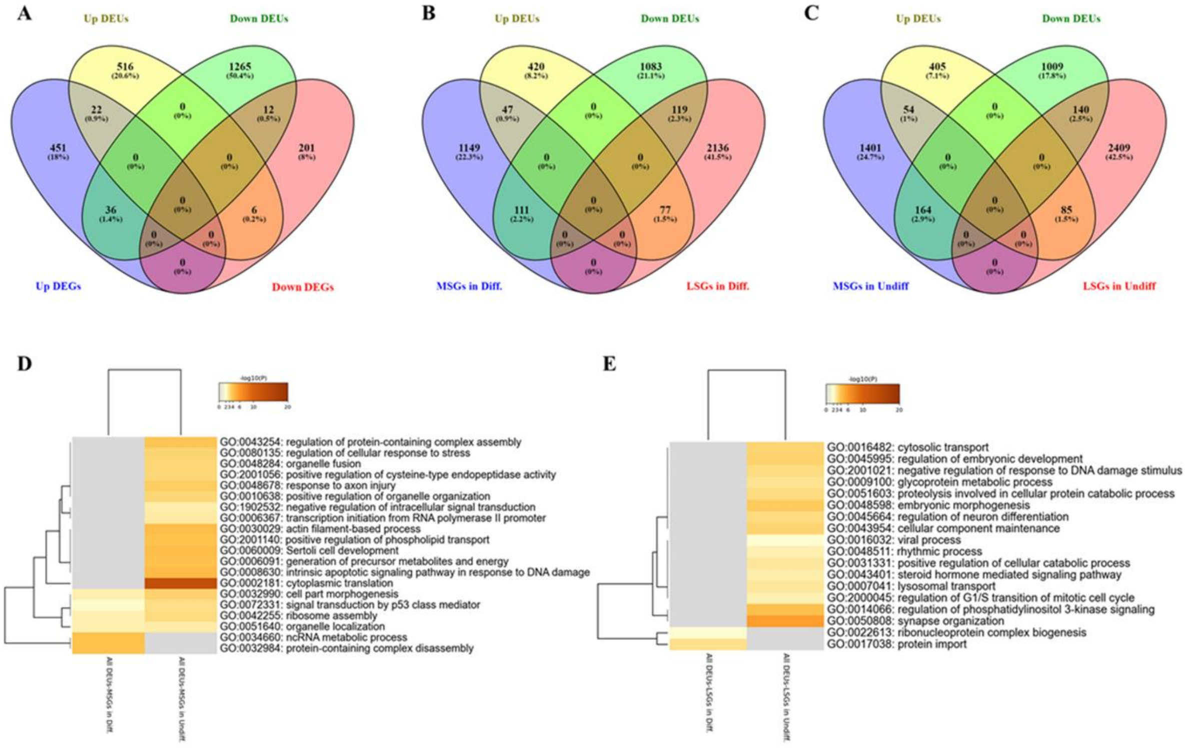

**Supplementary Figure 5** Comparative analysis between transcriptomics and differential alternative splicing events in differentiated and undifferentiated SH-SY5Y cells. **Related to Figures 2 and 5**. Venn diagram showing the ratio of overlapping genes among DEGs and each alternative splicing events SE (**A**), A5SS (**B**), A3SS (**C**), RI (**D**) and MXE (**E**), respectively.

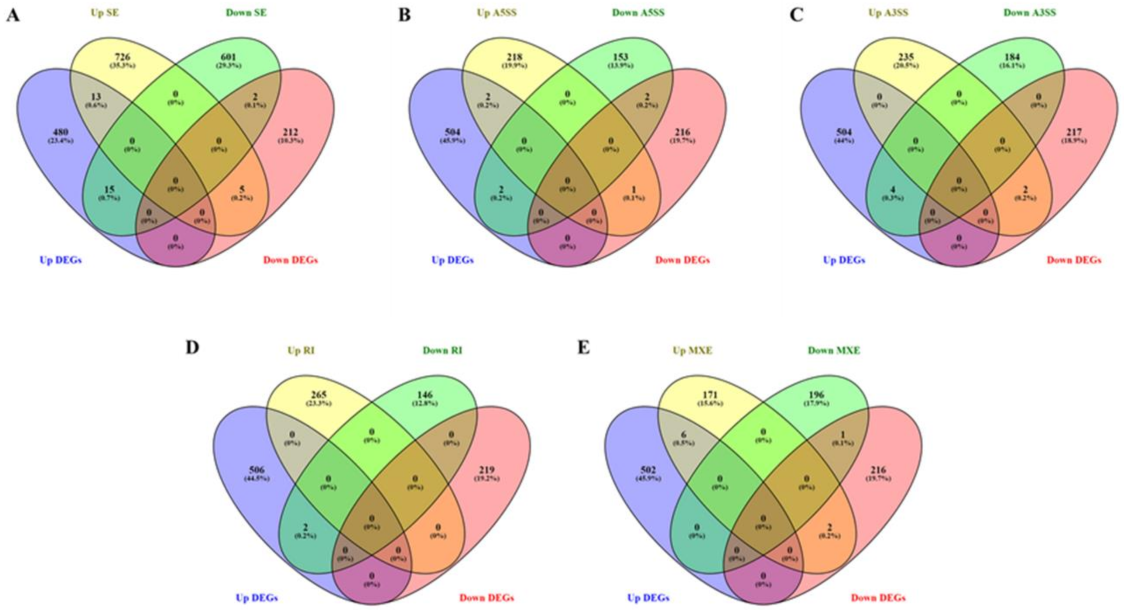

**Supplementary Figure 6** Comparative analysis between mRNA stability profiling and differential alternative splicing events in differentiated and undifferentiated SH-SY5Y cells. **Related to Figures 3 and 5.** Venn diagram showing the ratio of overlapping genes among MSGs and LSGs in Diff. and each alternative splicing events SE (**A**), A5SS (**C**), A3SS (**E**), RI (**G**) and MXE (**I**), respectively. Venn diagram showing the ratio of overlapping genes among MSGs and LSGs in Undiff. and each alternative splicing events SE (**B**), A5SS (**D**), A3SS (**F**), RI (**H**) and MXE (**J**), respectively.

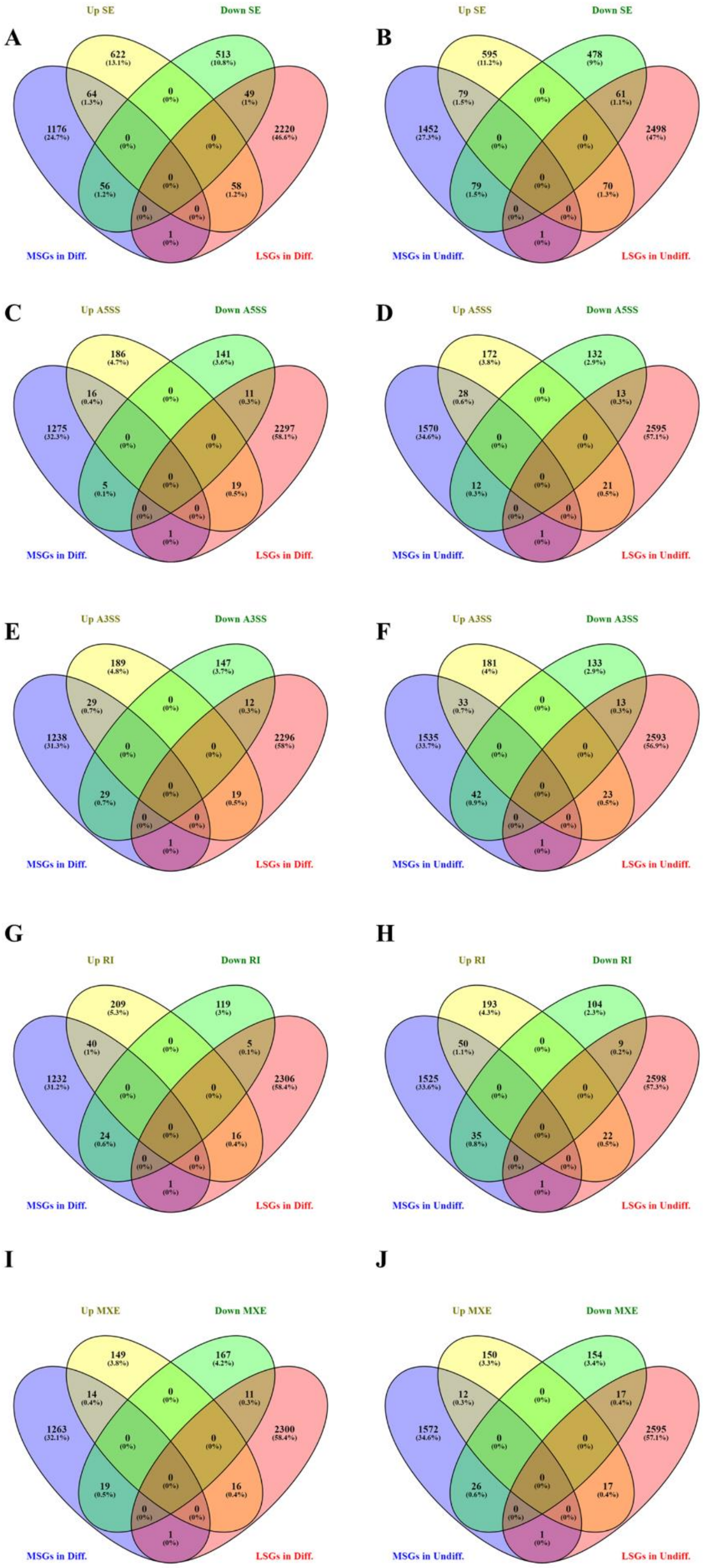
A
Up SE
Down SE
622
513
64
0
49
1176
0
0
2220
0
56
58
0
0
1
MSGs in Diff.
LSGs in Diff.
B
Up SE
Down SE
595
478
79
0
61
1452
0
0
2498
0
79
70
0
0
1
MSGs in Undiff.
LSGs in Undiff.
C
Up A5SS
Down A5SS
186
141
16
0
11
1275
0
0
2297
0
5
19
0
0
1
MSGs in Diff.
LSGs in Diff.
D
Up A5SS
Down A5SS
172
132
28
0
13
1570
0
0
2595
0
12
21
0
0
1
MSGs in Undiff.
LSGs in Undiff.
E
Up A3SS
Down A3SS
189
147
29
0
12
1238
0
0
2296
0
29
19
0
0
1
MSGs in Diff.
LSGs in Diff.
F
Up A3SS
Down A3SS
181
133
33
0
13
1535
0
0
2593
0
42
23
0
0
1
MSGs in Undiff.
LSGs in Undiff.
G
Up RI
Down RI
209
119
40
0
5
1232
0
0
2306
0
24
16
0
0
1
MSGs in Diff.
LSGs in Diff.
H
Up RI
Down RI
193
104
50
0
9
1525
0
0
2598
0
35
22
0
0
1
MSGs in Undiff.
LSGs in Undiff.
I
Up MXE
Down MXE
149
167
14
0
11
1263
0
0
2300
0
19
16
0
0
1
MSGs in Diff.
LSGs in Diff.
J
Up MXE
Down MXE
150
154
12
0
17
1572
0
0
2595
0
26
17
0
0
1
MSGs in Undiff.
LSGs in Undiff.

**Supplementary Figure 7** Identification of most significant enriched RNA binding proteins (RBPs) motifs for each alternative splicing events. **Related to Figure 6**. RNA motif maps of the most significant changed RBPs 9G8 (**A**) and RMB45 (**B**) in up- and down-regulated A5SS events respectively. RNA motif maps of the most significant changed RBPs LIN28A (**C**) and SRp20 (**D**) in up- and down-regulated A3SS events respectively. RNA motif maps of the most significant changed RBPs SRp20 (**E**) and RMB45 (**F**) in up- and down-regulated RI events respectively. RNA motif maps of the most significant changed BRUNOL5 (**G**) and RBM5 (**H**) in up- and down-regulated A5SS events respectively. The dotted lines indicate the significance of enrichment versus background in. The solid lines indicate the motif score of upregulated and downregulated genes compared to background genes.

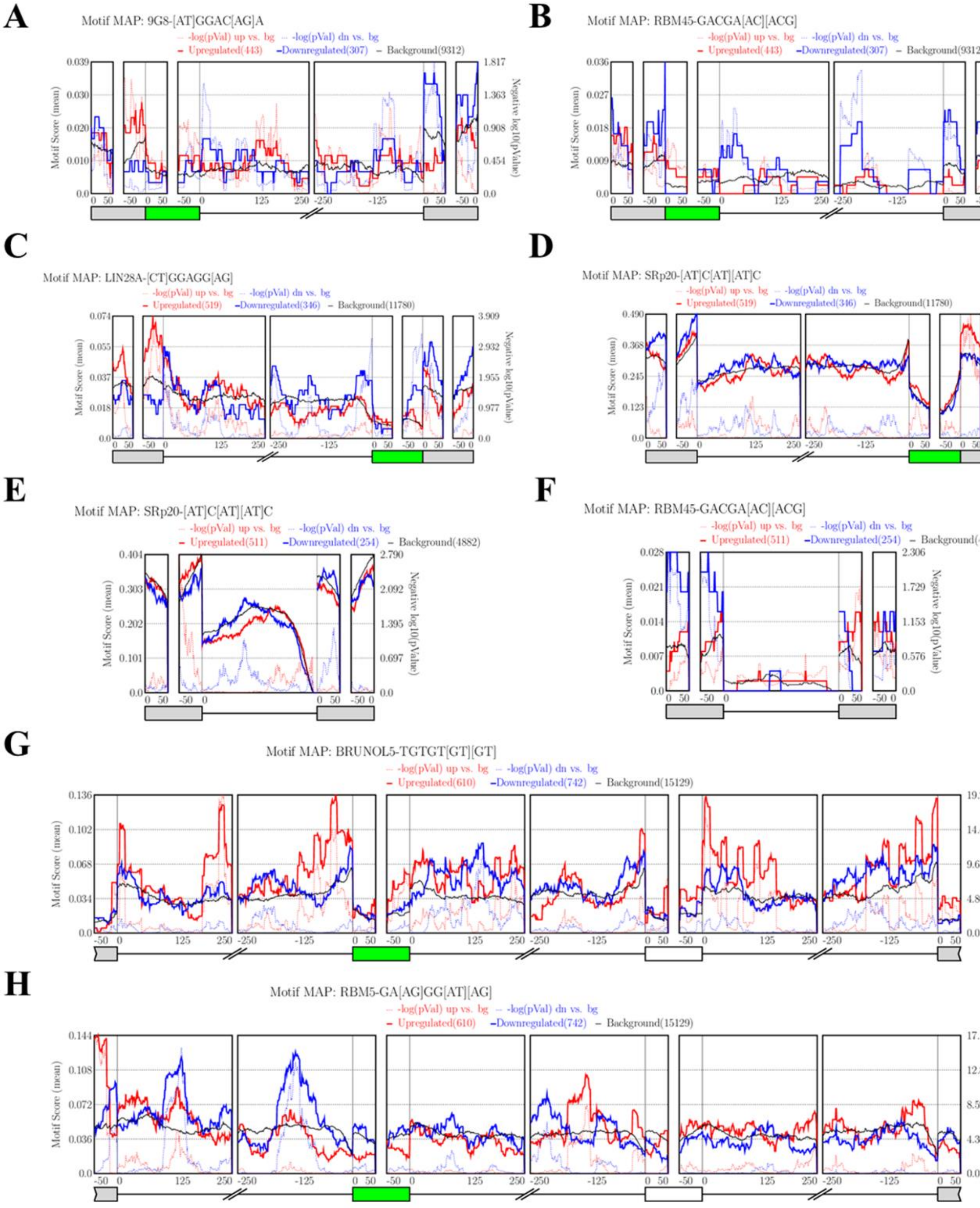
A
Motif MAP: 9G8-[AT]GGAC[AG]A
-log(pVal) up vs. bg
-log(pVal) dn vs. bg
Upregulated(443)
Downregulated(307)
Background(9312)
Motif Score (mean)
Negative log10(pValue)
B
Motif MAP: RBM45-GACGA[AC][ACG]
-log(pVal) up vs. bg
-log(pVal) dn vs. bg
Upregulated(443)
Downregulated(307)
Background(9312)
C
Motif MAP: LIN28A-[CT]GGAGG[AG]
-log(pVal) up vs. bg
-log(pVal) dn vs. bg
Upregulated(519)
Downregulated(346)
Background(11780)
D
Motif MAP: SRp20-[AT]C[AT][AT]C
-log(pVal) up vs. bg
-log(pVal) dn vs. bg
Upregulated(519)
Downregulated(346)
Background(11780)
E
Motif MAP: SRp20-[AT]C[AT][AT]C
-log(pVal) up vs. bg
-log(pVal) dn vs. bg
Upregulated(511)
Downregulated(254)
Background(4882)
F
Motif MAP: RBM45-GACGA[AC][ACG]
-log(pVal) up vs. bg
-log(pVal) dn vs. bg
Upregulated(511)
Downregulated(254)
Background(4882)
G
Motif MAP: BRUNOL5-TGTGT[GT][GT]
-log(pVal) up vs. bg
-log(pVal) dn vs. bg
Upregulated(610)
Downregulated(742)
Background(15129)
H
Motif MAP: RBM5-GA[AG]GG[AT][AG]
-log(pVal) up vs. bg
-log(pVal) dn vs. bg
Upregulated(610)
Downregulated(742)
Background(15129)

**Supplementary Figure 8** Comparative analysis between transcriptomics differentiated SH-SY5Y cells and CRISPRi/a screens on iPSC–derived neurons. **Related to Figures 2 and 7. A** Venn diagram on DEGs of differentiated SH-SY5Y and survival genes of CRISPRi screens on iPSC-derived neuron. **B** Venn diagram on DEGs of transcriptomic analysis on differentiated SH-SY5Y and survival genes of CRISPRa screens on iPSC-derived neuron.

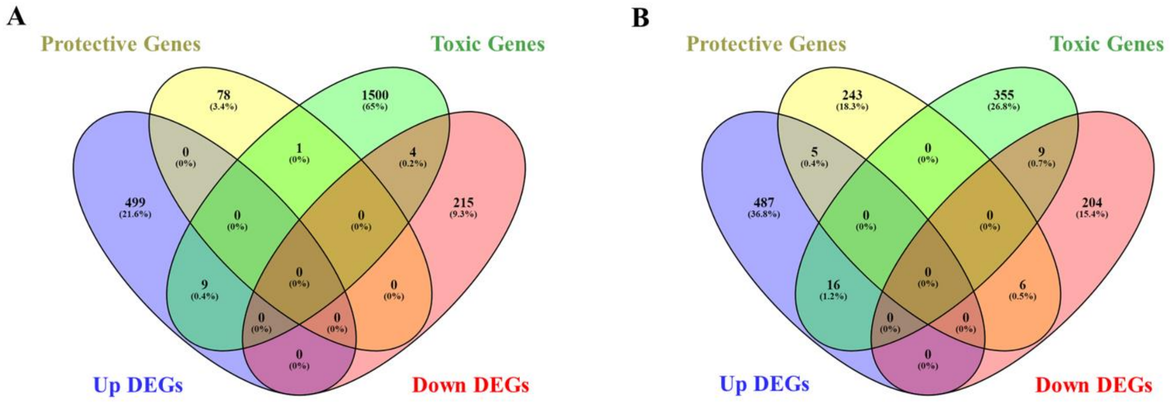
